# Evolutionary Transcriptome Analysis Based on Differentially Expressed (DE) Genes

**DOI:** 10.1101/2020.05.16.099804

**Authors:** Xun Gu

## Abstract

To address how gene regulation plays a key role in phenotypic innovations through high throughput transcriptomes, it is desirable to develop statistically-sound methods that enable researchers to study the pattern of transcriptome evolution. Most methods currently available are based on the Ornstein-Uhlenbeck (OU) model that considers the stabilizing selection as the baseline model of transcriptome evolution. In this paper, we developed a new evolutionary approach, based on the genome-wide *p*-value profile arising from statistical testing of differentially expressed (DE) genes between species. Our current approach is focused on the estimation of transcriptome distance between species. We first establish the relationship between the evolutionary model (the Markov-chain or Poisson model) and the proportion of null hypothesis (*u*_*0*_), which can be used to estimate the transcriptome distance. Further, we calculate the posterior probability of a gene being DE when a *p*-value is given, denoted by *Q*=*P*(*DE|p*), and develop a simple algorithm to estimate the transcriptome distance for any number of genes in the genome. Our compute simulations showed the statistical performance of these new methods are generally satisfactory.

## Introduction

With enormous amounts of transcriptome (RNA-seq) data from multiple tissues with diverse species available, it is the exciting time for evolutionary biologists to address how gene regulation plays a key role in phenotypic innovations (King and Wilson 1975; Lehner 2013) (e.g., Wang, et al. 2009; Brawand, et al. 2011; McCarthy, et al. 2012; Necsulea and Kaessmann 2014; Xu, et al. 2018; Cardoso-Moreira, et al. 2019). To help achieve this goal, development of statistically-sound methods that enable researchers to study the pattern of transcriptome evolution is desirable (Gu and Su 2007; Pereira, et al. 2009; Gu, et al. 2013; Gu 2016; Ruan, et al. 2016; Gu, et al. 2017; Sudmant, et al. 2015; Yang et al. 2018; Gu et al. 2019). We (Yang et al. 2019) recently developed an R-package, *TreeExp2* (*Tree*-dependent *Exp*ression analysis for short), which provides a suite of up-to-date statistical methods for phylogenetic analysis of transcriptomes. It should be noticed that most methods implemented in *TreeExp2* are based on the Ornstein-Uhlenbeck (OU) model that considers the stabilizing selection as the baseline model of transcriptome evolution (Hansen and Martins 1996; Lemos, et al. 2005), that is, stabilizing selection, which maintains the optima under the background of random mutations, dominates the transcriptome evolution (Bedford and Hartl 2009).

In a high-throughput transcriptome study, statistical detection of differentially expressed (DE) genes between two sets of samples is probably the most common practice in the transcriptome data analysis (Cheng et al. 2004; Efron 2004; Efron et al. 2001). Each statistical hypothesis testing (usually for each gene) results in a *p*-value summarizing the level of statistical significance of the observed differential gene expression. Xu et al. (2018) conducted an analysis to predict differentially expressed (DE) genes between the human and chimpanzee in several brain regions. Their study indicated that detection of DE genes between species is a valuable approach to understanding the pattern of transcriptome evolution.

In the past decades a wealth of statistical literatures were reported to deal with the multiple DE-test problem, i.e., many null hypotheses being tested simultaneously in a high throughput experimentation. Our goal is to formulate an evolutionary model based on the statistical framework primarily developed for the detection of DE genes (Benjamini and Hochberg 1995; Benjamini and Yekutieli 2001). In this paper we focus on the estimation of transcriptome distance between species, a basic parameter for understanding the regulatory evolution. After invoking the widely-used BUM (beta-uniform mixture) model for the genome-wide *p*-value distribution (Pounds and Morris 2003; Smyth 2004; Storey and Tibshirani 2003; Zhang and Grant 2004; Zhang 2011), we first study the relationship between the evolutionary model (the Markov-chain or Poisson model) and the proportion of null hypothesis (*u*_*0*_), which can be used to estimate the transcriptome distance. Further, we calculate the posterior probability of a gene being DE denoted by *Q*=*P*(*DE|p*) based on the BUM model, and develop a simple algorithm to estimate the transcriptome distance for any number of genes with special interests. While compute simulation is used to evaluate the statistical performance of these new methods, case studies demonstrate their potential applications. The differences between these DE-based methods and OU-based methods are discussed.

## Results

### Statistically modeling of *p*-values for differentially expressed genes

While *p*-values arising from the null hypothesis are distributed uniformly on the interval (0, 1), those arising from the alternative hypothesis follows a distribution that is generally modeled by a beta distribution whose pdf is given by

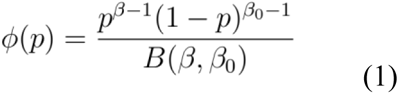

It follows that the distribution of a set of *p*-values can be written as a beta-uniform mixture model (BUM) consisting of a uniform (0, 1) component for the null hypothesis and a beta component for the alternative hypothesis, with the pdf given by

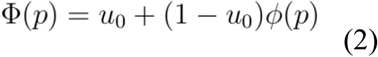

where *u*_*0*_ is the proportion of null hypothesis. The BUM model can be intuitively interpreted as follows. Under the null hypothesis, the *p*-values will have a uniform density corresponding to a flat horizontal line. Under the alternative hypothesis, the *p*-values will have a distribution that has high density for small *p*-values and the density will decrease as the *p*-values increase. The overall BUM distribution will be a mixture of values arising from the two hypotheses. Note that Eq.(1) involves two parameters, *β* and *β*_*0*_. With the restrictions of 0<*β*<1 and *β*_*0*_≥1, Eq.(2) provides a reasonable model for the distribution of *p*-values arising from high throughput genomics (Storey and Tibshirani 2003): it asymptotes at *p*=0 and monotonically decreases to its minimum of *u*_*0*_ at *p*=1.

### *u*_*0*_-Evolutionary distance of transcriptome

When the differentially expressed (DE) gene detection is applied to two species samples of the same tissue, one may speculate that the number of DE genes is small between closely-related species, whereas it is large between distantly-related species. An intriguing question is how it may relate to the distance of transcriptome evolution. As shown below, we will address this issue under two models: the Markov chain model, and the Poisson model.

#### The Markov-chain model

Suppose any gene in a given tissue has two states: unexpressed or expressed. Let π_1_ be the proportion of expressed genes, and π_0_=1-π_1_ be that of unexpressed genes. Let λ be the rate of transcriptome evolution. Under a two-state Markov-chain model, the gain rate and the loss rate of an expression status are given by π_1_λ and π_0_λ, respectively. It follows that the transition probabilities from state *i* to state *j* after the *t* time units, *P*_*ij*_(*t*), *i, j*=0 (unexpressed), or 1 (expressed), are given by *P*_*11*_(*t*)= π_1_+π_0_*e*^−*λt*^, and *P*_*10*_(*t*)= π_0_(1-*e*^−*λt*^), respectively (*t* is the speciation time), and similarly, we have *P*_*00*_(*t*)=π_0_+π_1_*e*^−*λt*^, and *P*_*01*_(*t*)= π_1_(1-*e*^−*λt*^). For any orthologous gene pair, the probability for any observed expression pattern *P*(*r*_*A*_, *r*_*B*_) can be calculated as follows

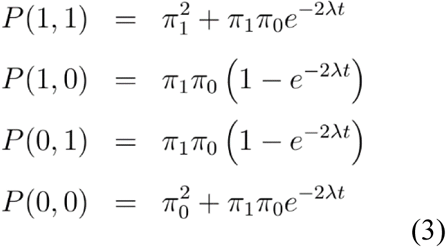

For instance, *P*(1, 1) is the probability that both genes are expressed; *P*(1, 0) is the probability that gene-1 is expressed but gene-2 is unexpressed; and so forth.

The Markov-chain model depicted by Eq.(3) provides an evolutionary interpretation for the BUM model of DE genes: the probability of a gene being no differentially expressed (null hypothesis) is given by *P*(1, 1)+*P*(0, 0), as predicted by *u*_*0*_, the proportion of null hypotheses. The relationship *u*_*0*_=*P*(1, 1)+*P*(0, 0) implies that

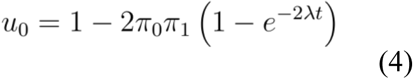

Apparently, a large proportion of non-DE genes is expected for closely-related speies (*u*_*0*_→1 as *t*→0), while it decreases for distantly-related specis, ultimately toward a saturation level of (1-2π_1_π_0_) as *t*→∞. Next, we define the distance of transcriptome evolution by *G*=(2π_1_π_0_)2λ*t*, the expected number of expression changes per gene per time unit. According to Eq.(4), one can estimate *G* by the following formula

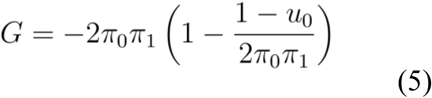

provided that *u*_*0*_, the proportion of null hypotheses, is estimated.

#### The Poisson model

Under the Poisson model, the probability of a gene being no differently expressed (non-DE) between two species is given by *e*^−2*λt*^. Hence, the *u*_*0*_-transcriptome distance defined by *G*=2*λt* can be simply estimated by the relationship *e*^−2*λt*^=*u*_*0*_, i.e.,

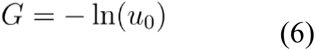

Usually, the Poisson distance given by Eq.(6) tends to underestimate the transcriptome distance. Yet, it is useful in practice in the case when π_1_ (or π_0_) is difficult to estimate, or when Eq.(5) is inapplicable, which may occur between two distantly-related species. At any rate, both Eq.(5) and Eq.(6) converge to 1-*u*_*0*_ for closely-related species.

### Estimation procedure of *u*_*0*_-transcriptome distance and case study

#### Non-parametric estimation of *u*_*0*_

Suppose we have multiple hypothesis tests with simultaneously *M p*-values. With some minor modifications, we implemented the algorithm developed by Zhang and Grant (2004) to estimate *u*_*0*_, the proportion of null hypotheses. First, the *M p*-values are first sorted in ascending order so that *p_1_≤p_2_≤p_3_≤…p_M-1_≤p_M_*. An empirical cumulative distribution of *p*-values is obtained by plotting *i/M* versus *p*_*i*_. Second, we connect each point (*i/M*, *p*_*i*_) on the empirical cumulative plot to the end point (1.0, 1.0) by a straight line, and calculates the slope of the line as

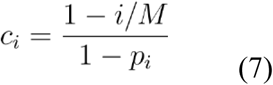

Then the median value of *c*_*i*_ for a given range of *p*_*i*_, *p*_*l*_≤*p*_*i*_≤*p*_*u*_, is taken as an estimate of *u*_*0*_,

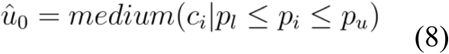

with *p*_*l*_=0.40, *p*_*u*_=0.95 as the default values of the method parameters.

#### Data availability

We downloaded published RNA-seq datasets of four primates, human, chimpanzee, gorilla and gibbon (Xu et al. 2108). Each species consists of eight brain areas, including cerebellum (CB), hippocampus (HIP), striatum (STR), anterior cingulate cortex (ACC), primary visual cortex (V1C), premotor cortex (PMC), ventrolateral prefrontal cortex (VPFC), DPFC (dorsolateral prefrontal cortex (DPFC). All primate species possess 2-6 biological replicates except that gibbon only has one for all samples. Among the cerebral region, the neocortical area includes ACC, V1C, PMC, VPFC and DPFC, whereas the subcortical area includes HIP and STR. For each RNA-seq sample, the median ratio method was applied to normalize the raw reads count data to make these expression values comparable cross species and brain areas. Following the original authors, a total of 27,991 expressed genes are used in the current study.

#### The u0-transcriptome distances between human and chimpanzee brain regions

For each of eight brain regions, we carried out the differentially expressed (DE) gene analyses between the human and chimpanzee, using three software packages, *DESeq2* (Love et al. 2014), *edgeR* (Robison et al. 2010) and *limma* (Ritchie et al. 2015). The genome-wide *p*-value profile for each method in each brain region was obtained. We then estimated *u*_*0*_, the proportion of null hypothesis, in each case by Eq.(7) and Eq.(8). As shown in Table 1, the estimate of *u*_*0*_ varies among methods as well as brain regions. The average *u*_*0*_ (over brain regions) is 0.88, 0.91.and 0.95 for *DESeq2*, *edgeR* and *limma*, respectively. The range of *u*_*0*_ by *DESeq2*, for instance, is from 0.825 (CB) to 0.914 (PMC).

**Table 1.** Transcriptome distances between the human and chimpanzee brain regions

| Methods | $u_0$-transcriptome distance <sup>b</sup> | | | | | | SOU distance <sup>d</sup> |
| --- | --- | --- | --- | --- | --- | --- | --- |
|  | *DEseq2* |  | edgeR |  | limma |  |  |
| Brain regions <sup>a</sup> | $u_0$ | $G$ | $u_0$ | $G$ | $u_0$ | $G$ | $D$ |
| CB | 0.825 | 0.2300 | 0.854 | 0.1813 | 0.881 | 0.1411 | 0.0899 |
| HIP | 0.850 | 0.1874 | 0.883 | 0.1381 | 0.961 | 0.0414 | 0.0974 |
| STR | 0.875 | 0.1502 | 0.917 | 0.0924 | 0.931 | 0.0760 | 0.0816 |
| ACC | 0.883 | 0.1380 | 0.893 | 0.1243 | 0.965 | 0.0367 | 0.0712 |
| V1C | 0.901 | 0.1136 | 0.936 | 0.0692 | 0.966 | 0.0353 | 0.0850 |
| PMC | 0.914 | 0.0961 | 0.962 | 0.0397 | 0.983 | 0.0177 | 0.0808 |
| VPFC | 0.905 | 0.1083 | 0.943 | 0.0616 | 0.968 | 0.0331 | 0.0653 |
| DPFC | 0.885 | 0.1347 | 0.934 | 0.0724 | 0.965 | 0.0363 | 0.0708 |
| Mean<br>±s.e. | 0.880<br>$\pm 0.010$ | 0.1448<br>$\pm 0.0147$ | 0.915<br>$\pm 0.012$ | 0.0974<br>$\pm 0.0156$ | 0.952<br>$\pm 0.011$ | 0.0522<br>$\pm 0.0131$ | 0.0803<br>$\pm 0.0035$ |

Notes. <sup>a</sup> brain regions: cerebellum (CB), hippocampus (HIP), striatum (STR), anterior cingulate cortex (ACC), primary visual cortex (V1C), premotor cortex (PMC), ventrolateral prefrontal cortex (VPFC), and dorsolateral prefrontal cortex (DPFC). <sup>b</sup> $u_0$ -transcriptome distance ( $G$ ) based on three DE detection software packages (DESeq2, edgeR and Limma). <sup>d</sup> The transcriptome distance based on the stationary OU model.

To estimate *u*_*0*_-transcriptome distance (*G*) under the Markov-chain model, we have to estimate π_1_ and π_0_. From ENSEMBL, the total number of human genes is roughly 39K, one can estimate π_1_=27991/39000=0.717 and π_0_=0.283. For the comparison, we estimated the transcriptome distance under the stationary OU model (Yang et al. 2019). Interestingly, for each brain region, the estimate of SOU transcriptome distance is somewhere among three *u*_*0*_-transcriptome distances (Fig.1). In other words, differences among statistical methods for DE detections may accounts for most estimation differences. Yet the average *u*_*0*_-distance is actually impressively consistent with the SOU-distance. As shown below, the new evolutionary framework may provide some insights on the long-term research topic on how transcriptome evolved between the human and chimpanzee (Enard et al. 2002; Gu and Gu 2003; Gilad et al. 2006; Nowick et al. 2009; Brawand et al. 2011; Xun et al. 2018).

**Fig.1.**
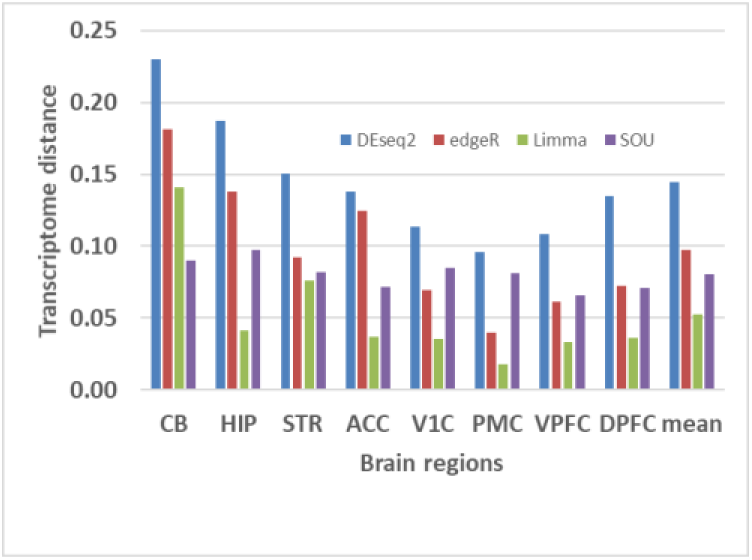
*u*_*0*_-Transcriptome distance between the human and chimpanzee brain regions, as well as the transcriptome distance based on stationary OU model (SOU for short)

### A Bayesian approach to estimating transcriptome distance

#### The posterior probability for a gene to be differentially expressed, given a p-value

Instead of the most common *p*-value assigned for each DE test, we anticipate that the posterior probability of a gene being differentially expressed, denoted by *P*(DE|*p*), may be useful for the analysis of transcriptome evolution (Pounds and Morris 2003; Smyth 2004; Storey and Tibshirani 2003). According to the BUM model of Eq.(2), it is straightforward to calculate *Q*=*P*(DE|*p*) by the Bayes rule, that is

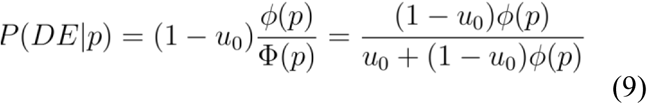

In the following analysis we use a special case of BUM with *β*_*0*_=1(Storey and Tibshirani 2003), in which *P*(DE|*p*) can be simplified as follows

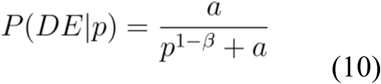

where *a*=(1-*u*_*0*_)*β*/*u*_*0*_. We implemented the following empirical Bayesian procedure: (*i*) estimate *u*_*0*_ by the nonparametric method of Eq.(8); and (*ii*) after treating the estimate of *u*_*0*_ as known, the maximum likelihood (ML) estimates of *β* in Eq.(2) can be obtained by fitting the beta-uniform mixture (BUM) distribution with *β*_*0*_=1 to the observed *p*-values (Pounds and Morris 2003; Storey and Tibshirani 2003).

#### Q-evolutionary distance of transcriptome

Suppose we have *M* hypothesis tests (*p*-values) at the genome level. Let *Q*_*k*_=*P*(DE|*p*_*k*_) be the posterior probability of the *k*-th gene being differentially expressed, *k*=1,…*M*. Intuitively, one may use the average *Q* given by

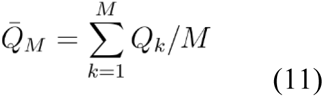

for empirically estimate of *q*=*P*(1, 0)+*P*(0, 1), the proportion of differentially expressed genes between species. Hence, under the Markov-chain model, the *Q*-transcriptome distance is given by

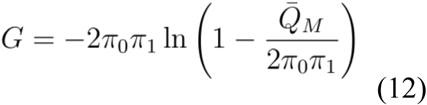

and that under the Posiison model given by

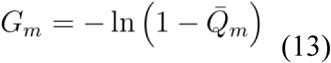

### Equivalence between the *u*_*0*_-based and *Q*-based transcriptome distances

An intriguing question is to what extent the *u*_*0*_-based and *Q*-based transcriptome distances may differ. We address this issue as follows. Since the posterior probability *P*(DE|*p*) conditional of *p* can be mathematically viewed as a function of *p*-value, one can calculate the conditional expectation of *P*(DE|*p*) with respect to Φ(*p*) under the BUM model, denoted by *E*[*P*(DE|*p*)]. From Eq.(9) one can show

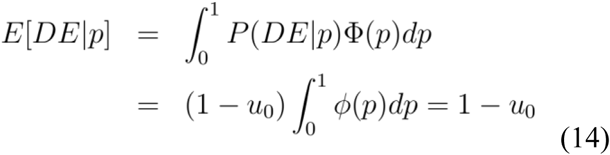

When the number of genes (*M*) is sufficently large, Eq.(14) implies that the average of *Q’*s given by Eq.(11) is expected to be

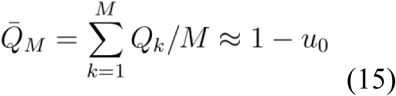

Therefore, the *u*_*0*_-based transcriptome distance and the *Q*-based transcriptome distances are equivalent by the means of empirical Bayesian: to calculate the posterior probability *Q*_*k*_=*P*(DE|*p*_*k*_), one may choose the prior *u*_*0*_ as the empirical estimate from the data.

### *Q*-based transcriptome distance for a small set of genes

Some important issues in evolutionary genomics require transcriptome distance analysis of a specific set (*m*) of genes; for instance, to evaluate a lineage-specifc fast-evolving pattern. However, when the number of genes (*m*) is small, estimation of *u*_*0*_ under the BUM model is notoriously difficult (Pounds and Morris 2003; Smyth 2004; Storey and Tibshirani 2003; Zhang and Grant 2004; Zhang 2011): our simulation study (see below) showed that estimation of *u*_*0*_ could be subjet to a large sampling variance as well as a large bias when *m* is around hundreds or less. We developed an algorithm that provides a practically feasible solution, which includes the folowing steps.

i. Estimation of *u*_*0*_ and *β* by the methods we developed above, based on genome-wide (*M* genes) *p*-values.
ii. In the case of Markov-chain model, π_1_ is estimated by the proportion of genes expressed in the transcriptome sample, and π_0_=1-π_1_.
iii. Suppose we have *m* genes under study (*m*<*M*). Calculate the posterior probability of each gene being DE when the *p*-value is given, denoted by *Q*_*k*_=*P*(DE|*p*_*k*_), *k*=1,…, *m*. And the average of posterior probabilities of being DE over *m* genes is given by

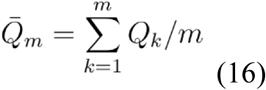
iv. Let *λ*_*m*_ be the evolutionary rate of transcriptome for those *m* genes. In analogy to Eq.(12), under the Markov-chain model, the transcriptome distance defined by *G*_*m*_=(2π_1_π_0_)*λ*_*m*_*t* can be estimated by

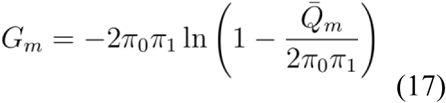

while under the Poisson model the transcriptome distance defined by *G*_*m*_=2*λ*_*m*_*t* is given by

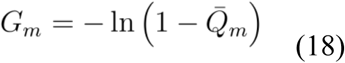

### Sampling variance of the transcriptome distance

While it is straightforward to estimate the transcriptome distance for any set of genes by Eq.(17) or Eq.(18), the question is how we can evaluate the statistical property of the distance estimation. There are two resources of estimation errors: the sample size, i.e., the number of genes under study (*m*), and the statistical uncertainty of high throughput experiments. Each gene *k* may have some contributions to this high throughput uncertainty, approximately measured by *Q*_*k*_(1-*Q*_*k*_) (Efron 2001); indeed, the contribution is the lowest (zero) if *Q*_*k*_=0 or 1 (no uncertainty), or the highest (0.25) if *Q*_*k*_=0.5 (maximum uncertainty).

We then developed an approximate method to estimate the sampling variance of transcriptome distance. By the delta method, we showed that the sampling variance of *Q*_*m*_ is approximately given by

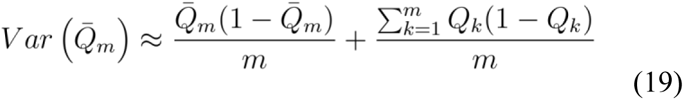

The first term on the right hand of Eq.(19) represents the sample size effect, which approaches to zero when the number of genes *m*→∞. The second term represents the variance due to high throughput uncertainty, which generally cannot vanish except for all *Q*_*k*_ is zero or one. It follows that the variance of *G*_*m*_ under the Markov-chain model is given by

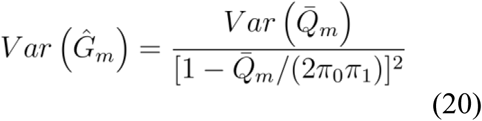

and that under the Poisson distance is given by

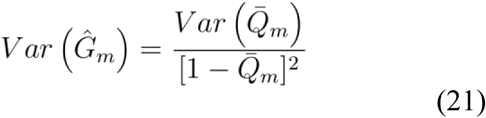

### Simulation studies

We carried out simulation studies to evaluate the statistical performance of *Q*-transcriptome distances for any number of genes, which can be concisely described as follows. *Theme-A.* (*i*) Under a two-species scenario of transcriptome evolution, simulate the expression patterns of *M* genes according to a specified two-state Markov-chain model, and then calculate *u*_*0*_. (*ii*) Simulate the BUM distribution under this *u*_*0*_ and a specified *β* for a *p*-value profile of *M* hypotheses. (*iii*) Estimate *u*_*0*_ and *β* based on those simulated *p*-values. *Theme-B*. Repeat the first two steps in Theme-A for pre-specified *m* genes, under a given Markov-chain model and BUM model. After those *m p*-values are simulated, calculate the posterior probability *Q*_*k*_ for each gene, using the estimates of *u*_*0*_ and *β* by step-(*iii*) in Theme-A. In both themes *A* and *B*, the simulation will be repeated for 1000 rounds and then evaluate the statistical performance in each simulation case.

Our main results are summarized in Table 2. (*i*) Estimations of both *u*_*0*_ and *β* are statistically reliable and asymptotically unbiased as long as *M*>500. (*ii*) Estimation of the *Q*-transcriptome distance is, in general, asymptotically unbiased. That is, the estimation bias tends to negligible as long as the number of genes (*m*) is sufficiently large. (*iii*) The estimation variance decreases with the increasing of the number (*m*) of genes, but cannot vanish ultimately, due the high throughput uncertainty. (*iv*) In the case when *m* is sufficiently large, say, *m*>1000, the value of *β* is the key-factor to determine the accuracy of distance estimation. And (*v*) the variance of distance estimation given by Eq.(20) or Eq.(21) can be used as a good proxy in practice to the variance of distance estimation.

**Table 2.**
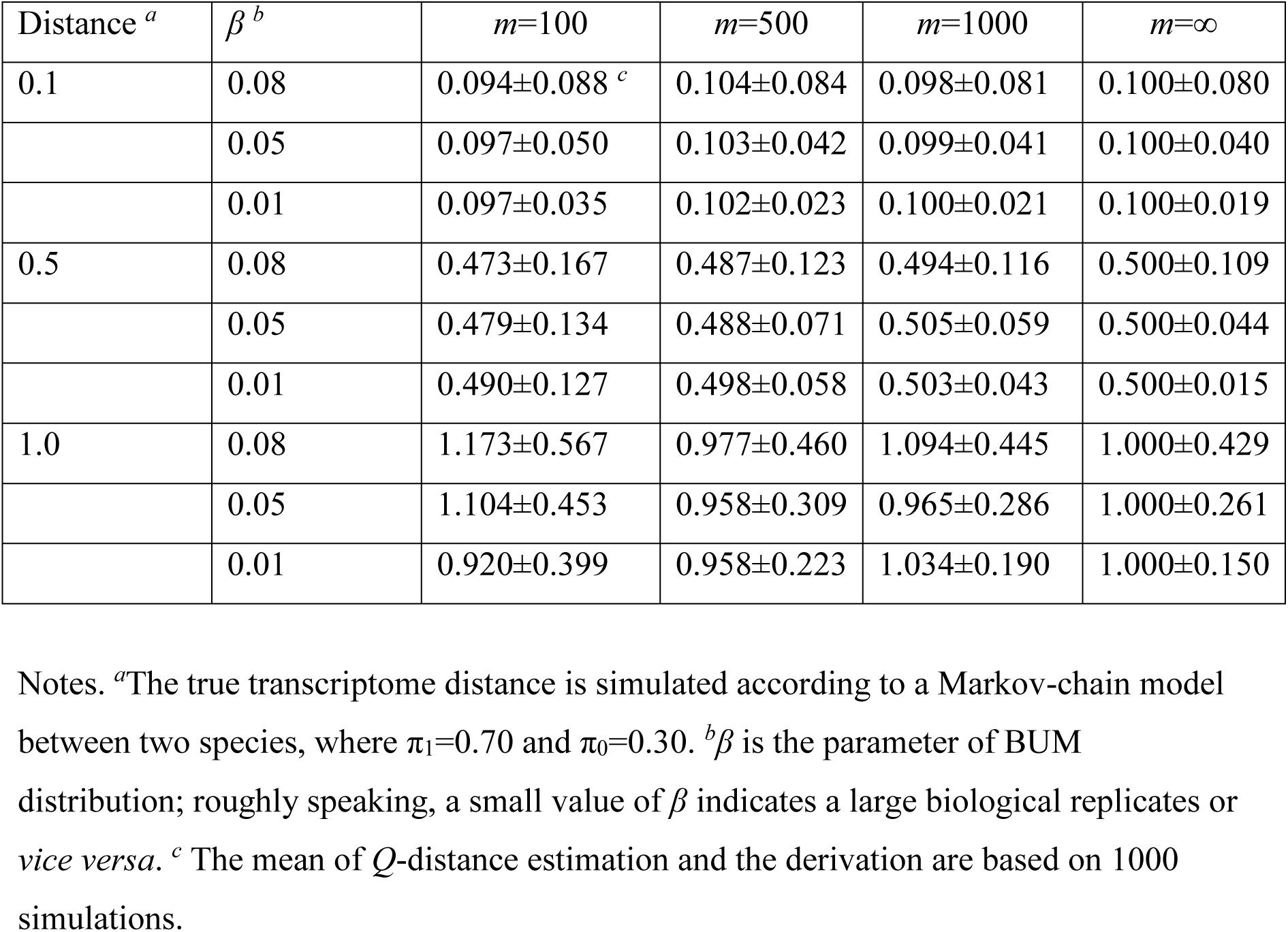
Simulation results of *Q*-transcriptome distances for given the number of genes (*m*).

| Distance <sup>a</sup> | $\beta$ <sup>b</sup> | $m=100$ | $m=500$ | $m=1000$ | $m=\infty$ |
| --- | --- | --- | --- | --- | --- |
| 0.1 | 0.08 | 0.094±0.088 <sup>c</sup> | 0.104±0.084 | 0.098±0.081 | 0.100±0.080 |
|  | 0.05 | 0.097±0.050 | 0.103±0.042 | 0.099±0.041 | 0.100±0.040 |
|  | 0.01 | 0.097±0.035 | 0.102±0.023 | 0.100±0.021 | 0.100±0.019 |
| 0.5 | 0.08 | 0.473±0.167 | 0.487±0.123 | 0.494±0.116 | 0.500±0.109 |
|  | 0.05 | 0.479±0.134 | 0.488±0.071 | 0.505±0.059 | 0.500±0.044 |
|  | 0.01 | 0.490±0.127 | 0.498±0.058 | 0.503±0.043 | 0.500±0.015 |
| 1.0 | 0.08 | 1.173±0.567 | 0.977±0.460 | 1.094±0.445 | 1.000±0.429 |
|  | 0.05 | 1.104±0.453 | 0.958±0.309 | 0.965±0.286 | 1.000±0.261 |
|  | 0.01 | 0.920±0.399 | 0.958±0.223 | 1.034±0.190 | 1.000±0.150 |

Notes. <sup>a</sup>The true transcriptome distance is simulated according to a Markov-chain model between two species, where $\pi_1=0.70$ and $\pi_0=0.30$ . <sup>b</sup> $\beta$ is the parameter of BUM distribution; roughly speaking, a small value of $\beta$ indicates a large biological replicates or *vice versa*. <sup>c</sup> The mean of $Q$ -distance estimation and the derivation are based on 1000 simulations.

## Discussion

### Independent assumption in the BUM model

The BUM model implicitly assumes that the multiple test statistics are independent (Pounds and Morris 2003; Smyth 2004; Storey and Tibshirani 2003; Zhang and Grant 2004; Zhang 2011). Because this independent assumption is likely unrealistic in the context of transcriptome experiments, estimation of *u*_*0*_ under the BUM would be biased. Indeed, estimating the proportion of null genes with nontrivial inter-gene dependence is beyond the scope of current study, which has been shown very difficult (Zhang 2011). While this independent assumption makes our new methods of transcriptome distance practically feasible, one should be cautious that it might not be adequate in some extreme cases.

### Underlying distribution of gene expressions

The new method for estimating the evolutionary distance of transcriptome are based on the BUM model of *p*-values. While the advantage is flexibility and wide applicability, the effect of the underlying gene-expression distribution on the accuracy of *u*_*0*_-estimation, as well as other parameters (*β* and *β*_*0*_), remains largely unknown. It would be interesting to see how the underlying distributions affect the accuracies of these u0-estimation methods, especially how robust these p-value based methods are when the actual distribution of gene expression departs from the assumed theoretical distribution. This would form the basis of a future investigation (Pounds and Morris 2003; Smyth 2004; Storey and Tibshirani 2003; Zhang and Grant 2004; Zhang 2011).

### Different methods for u0-estimation

The method here to estimate *u*_*0*_ is nonparametric, which does not depend on the specific form of statistical tests being used, as long as the *p*-values pertaining to the tests are obtained. Since there are many different methods proposed to estimate *u*_*0*_, it remains an open question to what extent the estimation of the evolutionary distance of transcriptome is sensitive to different methods used. In the future, we will carry extensive computer simulations to examine the relative performance of our transcriptome distance estimation to the different methods of *u*_*0*_-estimation (Pounds and Morris 2003; Smyth 2004; Storey and Tibshirani 2003; Zhang and Grant 2004; Zhang 2011).

### Effects of biological replicates

Both theoretical analysis and computer simulation has established an inverse relationship between the number of biological replicates (*R*) and the BUM parameter (*β*) (Zhang and Grant 2004; Zhang 2011). Our simulation study showed that the sampling variance of the estimated transcriptome distance may inflate considerably when *β*=0.08 or more. According to previous study, it implies that the new method tends to be statistically unreliable when the biological replicates is small, say, *R*=3 or less. Hence, it is a challenge for an evolutionary researcher facing the prospect of biological replicates in order to achieve a reasonably accuracy of distance estimation.

### Implementation

Under the *R*-environment, we implement a computational procedure to analyze the evolutionary pattern of transcriptome evolution based on *p*-values. At the current stage, the software is available upon request.

